# A minimal RNA ligand for potent RIG-I activation in living mice

**DOI:** 10.1101/178343

**Authors:** Melissa M. Linehan, Thayne H. Dickey, Emanuela S. Molinari, Megan E. Fitzgerald, Olga Potapova, Akiko Iwasaki, Anna M. Pyle

## Abstract

We have developed highly potent synthetic activators of the vertebrate immune system that specifically target the RIG-I receptor. When introduced into mice, a family of short, triphosphorylated Stem Loop RNAs (SLRs) induces a potent interferon response and the activation of specific genes essential for antiviral defense. Using RNAseq, we provide the first *in-vivo* genome-wide view of the expression networks that are initiated upon RIG-I activation. We observe that SLRs specifically induce type I interferons, subsets of interferon-stimulated genes (ISGs), and cellular remodeling factors. By contrast, poly(I:C), which binds and activates multiple RNA sensors, induces type III interferons and several unique ISGs. The short length (10-14 base pairs) and robust function of SLRs in mice demonstrate that RIG-I forms active signaling complexes without oligomerizing on RNA. These findings demonstrate that SLRs are potent therapeutic and investigative tools for targeted modulation of the innate immune system.

## Introduction

RIG-I is an innate immune sensor that plays a key role in recognizing and responding to infection by RNA viruses (*1, 2*). RIG-I is activated in conditions that introduce a terminal, double-stranded RNA molecule into the cell. These molecules can include viral genomes, replication intermediates, and any other species containing a stable RNA duplex that is terminated with a 5’-triphosphate or diphosphate group (*3, 4*). RIG-I activation by RNA leads to the induction of type I interferon (IFN) genes through the activation of its adaptor molecule, MAVS (*5*). Another member of the RIG-I-like receptors (RLRs), MDA5, binds to long stretches of double-stranded RNA and similarly triggers type I IFN production through MAVS (*6*). Type I IFNs induce hundreds of genes, collectively known as IFN-stimulated genes (ISGs), which have a variety of antiviral effector functions (*7, 8*). Both the RNA duplex and 5’-triphosphate moieties are important for specific, high affinity binding and signaling by RIG-I (*9*-*11*), and through a series of recent crystal structures and functional studies of RNA recognition by the RIG-I receptor, the molecular basis for these effects has been elucidated (*3, 12*).

When appropriately delivered and modulated, RIG-I agonists would be promising tools for application in immuno-oncology (*13*-*16*), antiviral prophylaxis (*17*-*20*), and vaccine adjuvant development (*21*). However, all of these applications require a specific and potent RIG-I ligand that is functional *in vivo*. Previous work has suggested that RIG-I function can be controlled and exploited pharmacologically through stimulation with small, well-defined RNA ligands that are no larger than other therapeutically administered oligonucleotides (*9, 10, 16*). Structural studies, quantitative biochemical work, cell-based assays and imaging studies have all established that RIG-I is an “RNA end-capper” that encircles a 10-base-pair RNA duplex as a monomer and forms a network of specific interactions with the terminal base-pair and the 5’-triphosphate (*3, 10*-*12, 22*-*27*). RIG-I binding to short dsRNA is sufficient to trigger MAVS activation, and this process is enhanced by K63-ubiquitin, which promotes multimerization of the RIG-I CARD domains (*28*-*30*). On longer dsRNA (>40bp), RIG-I forms aggregated filaments *in vitro*, and signaling in cell culture is less dependent upon K63-ubiquitin (*29, 31, 32*). However, we do not know if RIG-I oligomerization on RNA is necessary for signaling *in vivo*. We sought to determine the ability of a minimal RIG-I ligand to signal *in vivo* and to monitor the effects on downstream gene expression in a living animal, providing the foundation for development of an RNA therapeutic.

To this end, we designed a set of <u>S</u>temloop <u>R</u>NA (SLR) molecules that present a single duplex terminus, and therefore bind only one RIG-I molecule. The opposite end of the duplex is blocked with a stable RNA tetraloop, thereby ensuring that RIG-I binds the triphosphorylated duplex terminus in a single, structurally-defined orientation. SLRs can be visualized as a short cord with a knot at one end that blocks protein binding. When RIG-I is constrained in this way, a robust interferon response is observed in mammalian cells. Here we compare selective RIG-I activation by SLRs to that of longer RNA molecules that have been used in previous studies of RIG-I function. We then introduce SLRs into mice and observe potent, MAVS-dependent IFN induction, thereby establishing the physical mechanism for RNA-stimulated RIG-I activation in animals and providing a powerful set of synthetic RIG-I agonists that can be applied as probes, mechanistic tools and pharmacological agents.

## Results

### SLRs are an optimal design for RIG-I recognition

Unlike other RNA molecules that have been developed as RIG-I activators, complexes between SLRs and RIG-I molecules have been characterized crystallographically (*3, 10, 23*), making it possible to visualize and optimize the molecular interaction networks that stabilize active RIG-I/RNA complexes. Relative to two-piece duplexes, the stemloop design provides simplicity, structural stability and resistance to nucleases, while presenting a single duplex terminus that fits precisely into the RNA binding pocket of RIG-I (Supp Fig. 1, 2).

In order to examine the relative potency of SLRs and compare them with larger, more complex ligands, we employed a well-established cell-based reporter assay in which IFN induction is monitored upon transfection with RNA (*22*). We tested the potency of two SLRs that vary in duplex length: SLR10 (containing 10 base pairs) and SLR14 (containing 14 base pairs). Recent studies indicate that RNAs bearing a diphosphate at the 5’ terminus are bona fide ligands for RIG-I (*33*). Thus, we compared the SLRs with a corresponding diphosphorylated stemloop (pp-SLR14), as well as stemloops lacking 5’-phosphate moieties (OH-SLR10 and OH-SLR14). We also included another control RNA, which is a 5’-triphosphorylated RNA that is the same length as SLR10, but is single-stranded and non structural (ppp-NS). In parallel, we examined the IFN response of a 19-base pair two-piece duplex that is commercially marketed as a RIG-I ligand (19mer dsRNA), and various other two-piece duplexes that have been utilized for previous structure/function studies (Fig. 1A and Supp. Fig. 1). In addition, all of the duplexes are compared with poly(I:C), which is polydisperse and of unknown structure.

**Figure 1.**
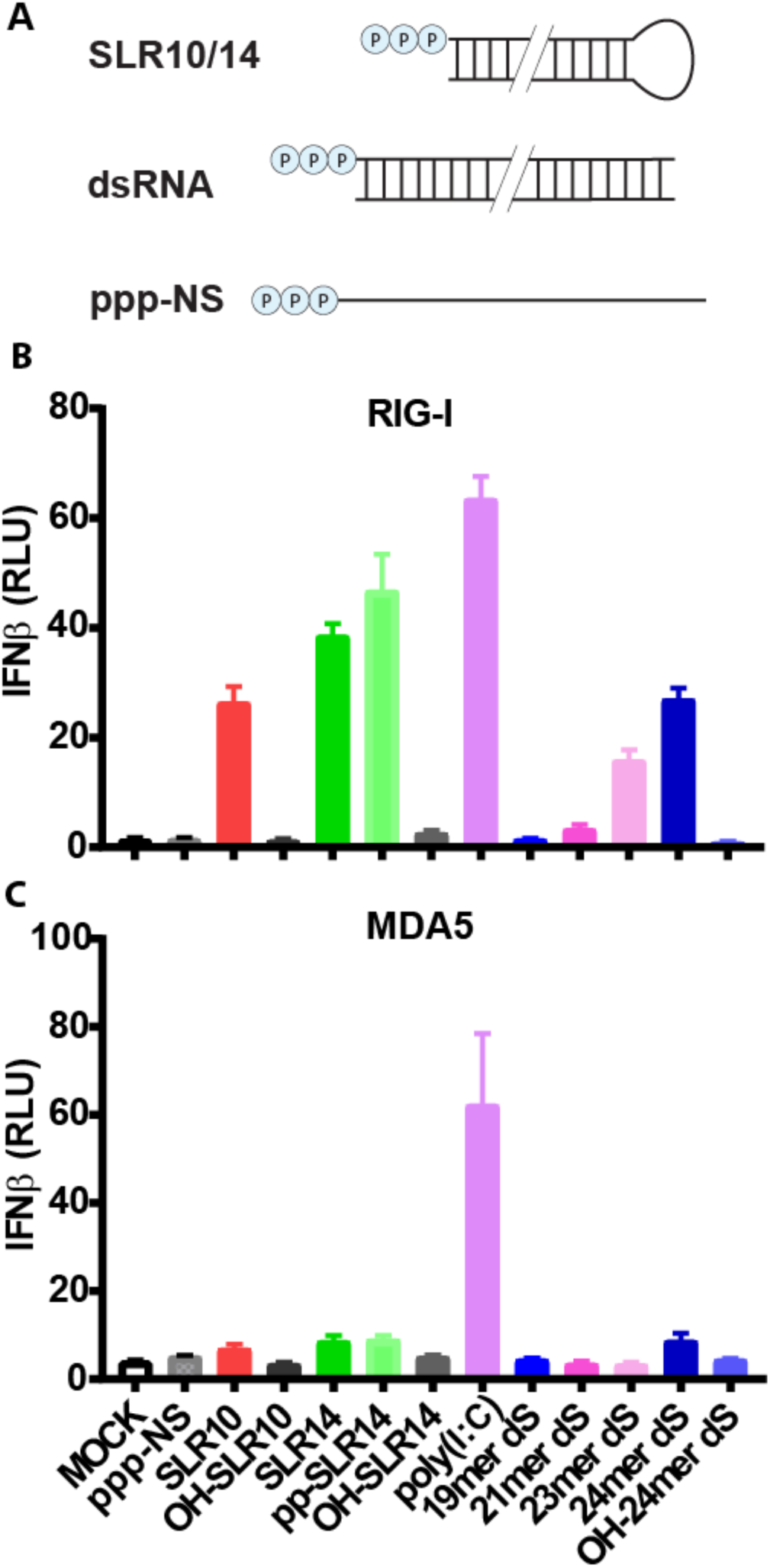
SLRs are optimally recognized by RIG-I. SLRs were designed to fold stably into a minimal RIG-I ligand containing 10 or 14 basepairs and a triphosphorylated 5’ terminus. SLRs were compared against other reported double-stranded RIG-I ligands (dsRNA) 19-24 basepairs in length and control RNAs lacking structure (ppp-NS) or 5’-triphosphates (OH-) (**A**). HEK293T cells lacking endogenous RIG-I and MDA5 were transfected with plasmids expressing RIG-I (**B**) or MDA5 (**C**) and a luciferase reporter under the control of an IFNβ promoter. Cells were then stimulated by various RNAs and luciferase production was measured as a proxy for IFNβ response. Poly(I:C) stimulates both receptors while SLRs are specific for RIG-I. SLR response is dependent upon the di- or tri-phosphate moiety and stimulates RIG-I as well, or better than other reported RIG-I ligands.

We observe that SLR10, SLR14 and pp-SLR14 are potent activators of RIG-I, inducing high levels of type I interferons (IFNs) in the cell-based assay (Fig. 1B). Relative to the 14-base pair duplexes, SLR10 displays somewhat reduced activity, but this trend is not universal (vide infra). As expected, the IFN response requires both of the determinants that are required for RIG-I activation: a 5’-di or -triphosphate and a duplex RNA structure, as OH-SLR10, OH-SLR14 and ppp-NS ligands failed to stimulate a significant level of IFN induction (Fig. 1B).

Notably, we failed to detect a significant IFN response with the RNA duplex (19mer dsRNA) that is being marketed and used as a specific RIG-I ligand (Supp. Fig. 1) (*32, 34*-*36*). We observed a small response from two RNA duplexes that have been utilized as RIG-I ligands in previous studies (*9*) (21mer dsRNA, 23mer dsRNA), and a moderate response from a double-stranded RNA that we specifically designed to have a high thermodynamic stability relative to previously published duplex designs (24mer dsRNA) (Fig. 1B). The IFN responses of the dsRNA ligands track closely with their relative thermodynamic stability (Supp Fig. 1), establishing that the efficacy of RIG-I ligands depends on a stable terminal duplex structure, which is enforced by the stemloop design in the SLRs. In keeping with previous findings, we observe a strong response from poly(I:C) (Fig. 1B). Taken together, these findings confirm previous work demonstrating the potency of SLRs in cell culture (*10, 37, 38*), and benchmark SLR potency relative to other putative RIG-I ligands.

### SLRs stimulate a potent IFN response in animals

In order to examine the ability of SLRs to induce type I IFN responses *in vivo*, we used a well established RNA delivery method to introduce RNA ligands intravenously in living mice (*17*). Wild type C57BL/6 mice were injected intravenously (IV) with RNA ligands complexed to polyethylenimine (PEI), and five hours later, IFN-α in mouse sera was measured by ELISA. A dose/response study of SLR10 showed an optimal response at 25 μg of RNA (Supp. Fig. 3). An RNA stemloop lacking a 5’-triphosphate (OH-SLR10) failed to induce IFN-α at any concentration (Supp. Fig. 3).

We then examined the IFN-α response to SLR10 molecules that were either synthesized (synth SLR10) or transcribed *in vitro* (trans SLR10) at the optimal RNA dose. We observed elevated levels of IFN-α after injection of synthetic SLR10 compared to transcribed SLR10 (Figure 2A). These results indicated that chemically synthesized SLR10, likely due to the enrichment in complete product, is more potent than transcribed RNA. Therefore, from here on, we exclusively use SLRs and dsRNA that are synthesized in house.

**Figure 2.**
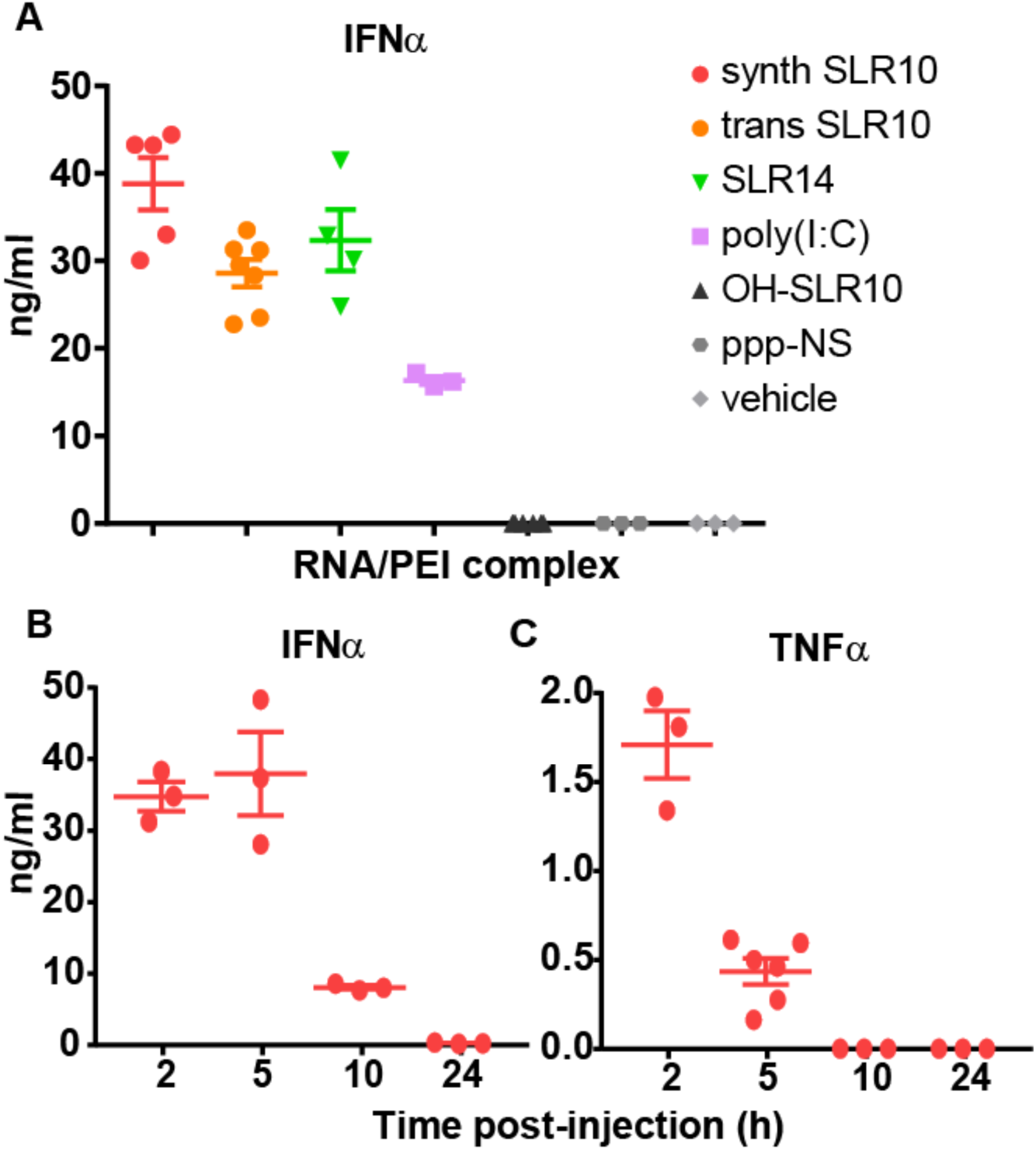
SLR injection induces robust type I IFN responses *in vivo*. C57BL/6 mice were injected intravenously (i.v.) with 25ug of SLR10, SLR14, OH-SLR10, poly(I:C), non-structural (ppp-NS), or vehicle control complexed with *in vivo*-jetPEI^®^ and sera were collected 5 hours later (**A**). C57BL/6 mice were injected i.v. with 25ug of synthesized SLR10 and sera were collected at the indicated time points (**B**, **C**). Synthetic RNA was used for all experiments except in (**A**) where *in vitro* transcribed SLR (trans SLR) was used for comparison. The concentrations of IFNα and TNFα were measured by ELISA. Synthetic SLR10 was superior to transcribed SLR10 in inducing serum IFNα.

We examined the extent of the IFN response induced by the various RNA ligands. Systemic IFN-α response was robustly induced by SLR10 and SLR14 (Figure 2A). In contrast, no induction of IFN-α was observed after OH-SLR10 or ppp-NS injection *in vivo*. The widely utilized ligand poly(I:C) induced IFN-α, but to a considerably lower degree (Figure 2A), thereby providing a benchmark for the relative *in vivo* activity of the SLR10 and SLR14 ligands.

Next, we analyzed the time course of IFN-α and TNF-α secretion in response to *in vivo* injected SLR10. By 2 hours post IV injection, SLR10 elicited robust IFN-α and TNF-α responses (Figure 2B, C). By 5 hours post injection, SLR10 sustained higher levels of IFN-α, which dropped off by 10 hours post-injection and were undetectable after 24 hours. In contrast, TNF-α levels peaked at 2 hours and were undetectable by 10 hours post-injection. These data indicate that SLRs induce rapid, robust and transient levels of IFN-α and TNF-α response in mice.

### Diphosphorylated stemloop RNAs are also potent inducers of type I IFN *in vivo*

To evaluate the ability of diphosphorylated RNA to trigger IFN responses *in vivo*, we injected 25 μg of pp-SLR10, OH-SLR10 or SLR10 into mice as described above, collecting the spleen after 5 hours of induction, and measuring the IFN-α mRNA by RT-qPCR. Both pp-SLR10 and SLR10 induced comparable levels of IFN-α mRNA in the spleen (Figure 3A). As expected, the non-phosphorylated OH-SLR10 induced minimal levels of IFN-α mRNA. In a separate experiment, we observed comparable ability of pp-SLR14 and SLR14 to induce IFN-α expression in the spleen (Figure 3B). Consistent with our *in vitro* (Figure 1) and *in vivo* analyses (Figure 2), both ppp-SLRs and pp-SLRs induced more robust IFN-α mRNA than poly(I:C) (Figure 3B). These results indicate that both in the serum (IFN-α protein; Figure 2) and spleen (IFN-α mRNA; Figure 3B), SLR and pp-SLR induce robust and comparable IFN-α expression.

**Figure 3.**
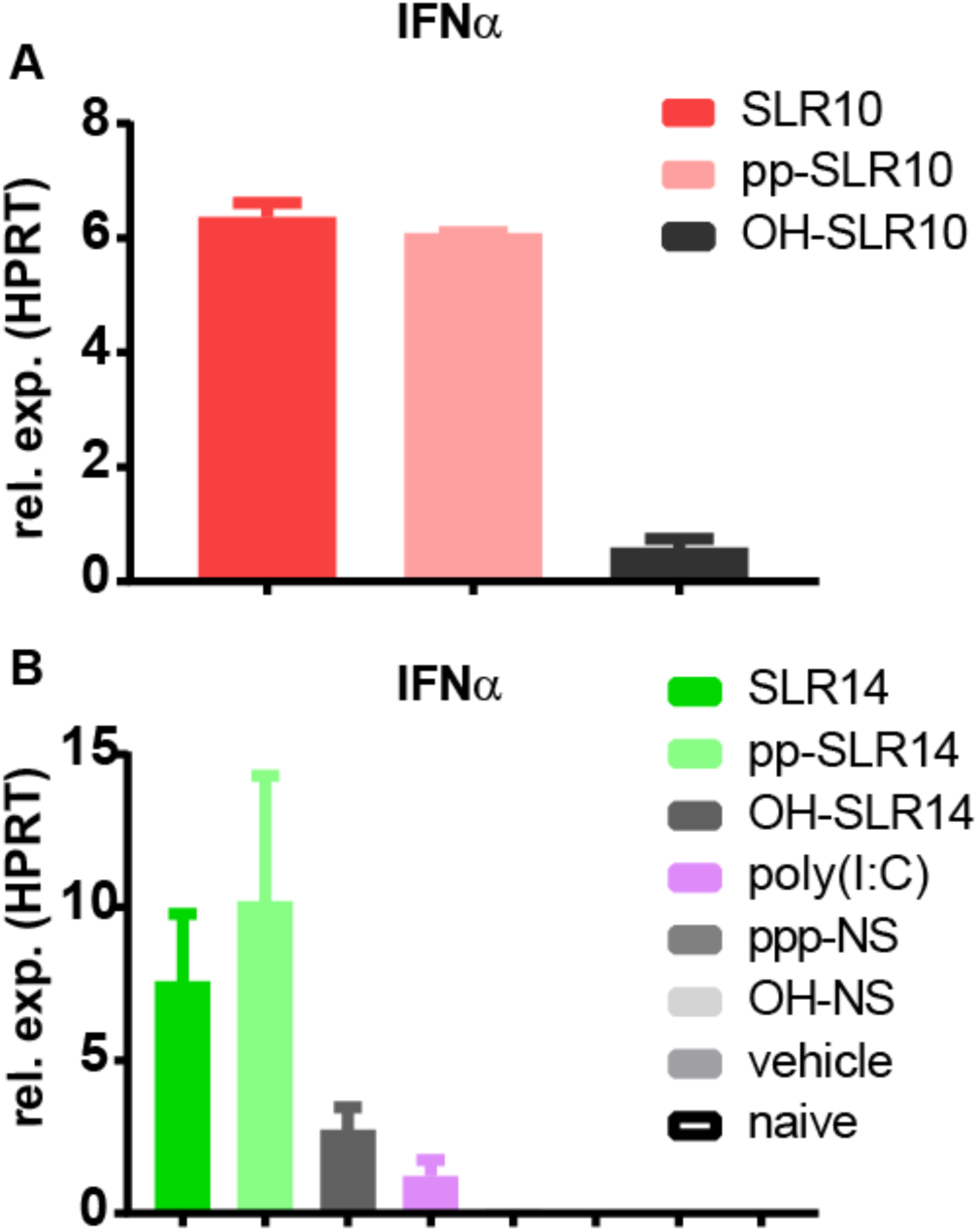
Diphosphorylated SLRs induce IFNα *in vivo*. The diphosphorylated counterparts to SLR10 (**A**) and SLR14 (**B**) were injected i.v. into C57BL/6 mice and 5 hours later, RNA was collected from spleens to measure IFNα by RT-qPCR. Both pp-SLR10 and pp-SLR14 induced comparable IFNα transcript levels as SLR10 and SLR14.

### Type I IFN induction by SLRs is RIG-I specific

Based on structural work and biochemical data (*12*), SLRs were designed to be a ligand that is specific for RIG-I, as other PRRs recognize nucleic acids with different types of features, such as extended length (MDA-5), or single-stranded character (TLR7). However, it was important to test the comparative PRR specificity of SLR ligands using a diversified set of experiments. Using the IFN-β promoter-luciferase reporter system described above, we found that both SLR10 and pp-SLR10 activate the IFN-β promoter when RIG-I is expressed, but not when MDA5 is expressed (Figure 1C). By contrast, poly(I:C) induced IFN-β promoter activation in both RIG-I and MDA5-expressing reporter cells, consistent with the fact that poly(I:C) contains recognition determinants important for both types of receptors (duplex termini and long RNA duplex regions).

To evaluate RIG-I specificity, we used siRNAs to knock down RIG-I expression in A549 lung epithelial cells. We then challenged the cells with SLRs and evaluated effects on IFN-β expression (Fig. 4A). Because knockdown of RIG-I was efficient in these experiments, it was possible to sensitively monitor the influence of challenge ligands on gene expression. Unlike their stimulatory behavior in WT cells, SLRs failed to induce IFN-β in the RIG-I KD cells (Fig. 4A). In addition, RIG-I KD resulted in diminished expression of ISGs, including *Rsad2* (viperin), and *Mx1* (Fig. 4A).These experiments show that SLRs are specifically and functionally recognized by RIG-I in human cells.

**Figure 4.**
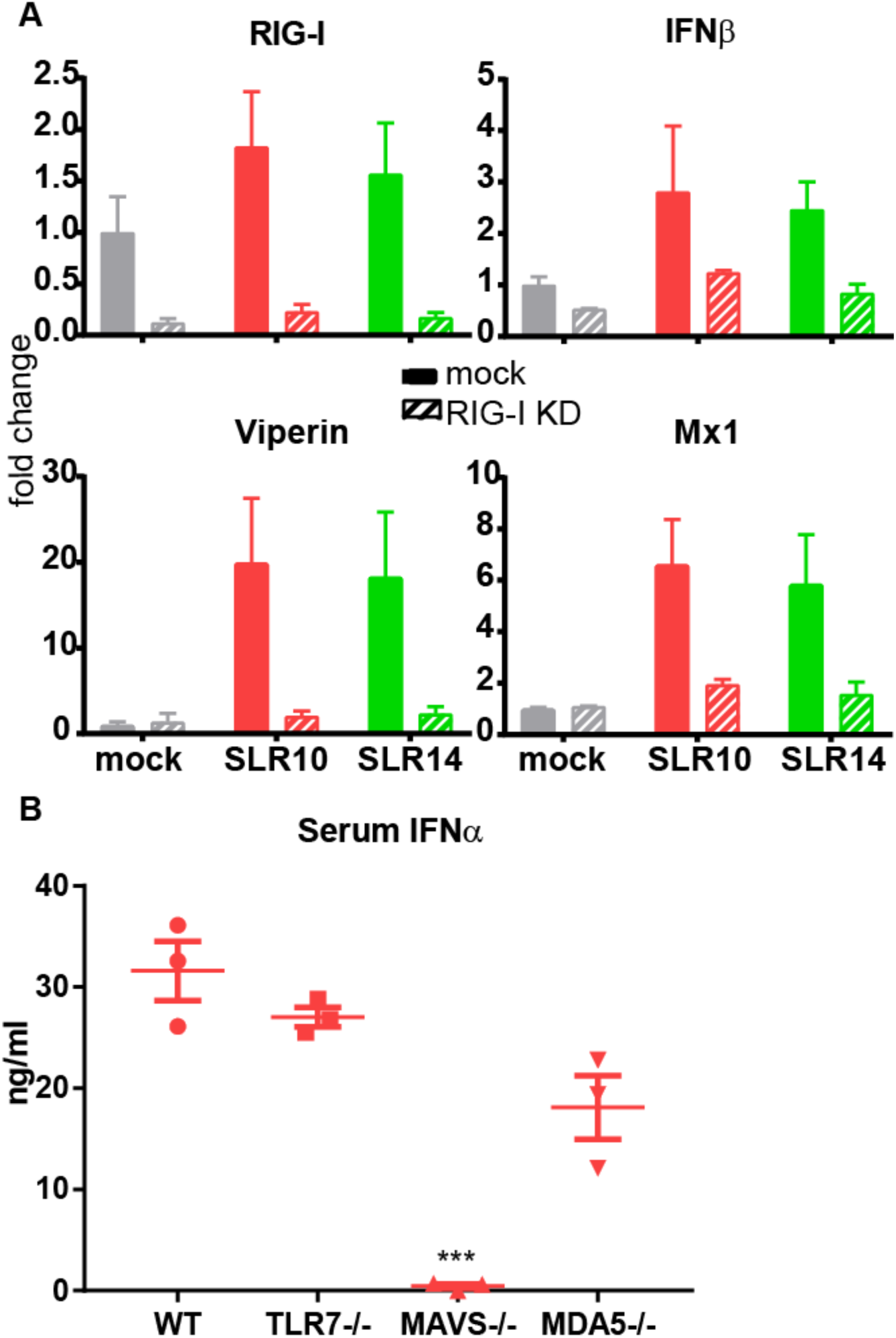
SLRs are specifically recognized by RIG-I. A549 human lung epithelial cells were transfected with mock or RIG-I targeting siRNAs, stimulated with SLR10 or SLR14 and assayed by RT-qPCR for knockdown efficiency and induction of IFNβ, Viperin and Mx1 (**A**). Reduction of RIG-I expression resulted in reduced IFN and ISG production. Mice lacking TLR7, MAVS, and MDA5 were injected i.v. with SLR10 and 5 hours later, sera were collected to measure IFNα. Asterisks indicate significant difference from WT mouse control (\*\*\**P <* 0.0005) (**B**).

To examine RIG-I specificity in a living animal, and to evaluate the potential contribution of other RNA sensors and signaling adaptors during IFN induction by SLRs, we introduced 25 μg of SLR10 i.v. into WT, *Tlr7*^-/-^, *Mavs*^-/-^ or *Mda5*^-/-^ mice. We were unable to examine the RIG-I deficient mice due to embryonic lethality (*39*). We collected sera at 5 hours and IFN-α levels were assessed by ELISA. These results demonstrate that MAVS, but not TLR7 or MDA5, is critical for IFN-α secretion following SLR injection (Figure 4B). The fact that SLRs require MAVS, but not MDA5, for IFN-α induction (Fig. 4B), and the fact that RIG-I knockdown eliminates this response in human cells (Fig. 4A) indicates that RIG-I is the receptor for IFN induction by SLRs.

### Specific activation of RIG-I by SLR and poly(I:C) reveals a distinct pattern of gene expression

Given the potential for SLR and other RNA agonists for clinical applications, it was of interest to determine the global gene expression profiles following injection of SLRs and other activating RNA ligands. In particular, it was important to explore the landscape of expressed genes in an unbiased manner so that we might be able to identify new processes involved in RIG-I induction. We therefore carried out RNAseq analysis of expressed genes in the mouse spleen 3 hours after injection of 25 μg of SLRs or poly(I:C). To our knowledge, this is the first time that the profile of RNA-induced gene expression has ever been determined in an animal.

Broadly speaking, we observe that SLR14 and SLR10, showed similar gene expression profiles (Supp. Fig. 4). To a first approximation, we can therefore use SLR14 as a proxy for short, multiphosphorylated RNA duplex ligands and compare them with RNAs that elicit a different profile of expressed genes, such as poly(I:C). We first measured the differentially expressed genes (DEGs) of SLR14 relative to vehicle, and compared that with the DEGs of poly(I:C) relative to vehicle. By comparing the profiles, we can assess the relative impact of poly(I:C) and SLR14 on patterns of gene expression.

Both SLR14 and poly(I:C) induced a shared set of genes involved in antiviral response and innate immunity, including *Ifit2, Oas3, Csf1, Tlr2, Tlr9, Nod2* and *Ddx58* (RIG-I) (Supp. Figure 4 and Supp. table 1). In addition, genes involved in T cell activation and effector functions (*Cd86, Tnf, Ifng, Il1b, Gzmb*) were elevated in response to both SLR14 and poly(I:C) (Supp. Table 1). Both SLR14 and poly(I:C) induced a marked downregulation in the expression of genes that belong in a variety of biological pathways, suggesting a shared pathway for co-opting basal cellular function upon induction. Many of these genes encode membrane proteins that are associated with processes such as ion transport, formation of cell junctions, and Wnt signaling (Supp. Figure 4 and Supp. Table 1). In addition, many myeloid expressed C-type lectins are downregulated, possibly indicating the loss of dendritic cell subsets from the spleen.

In contrast, the two types of ligands have differential effects on other gene families (Fig. 5 an SUPP.Table 2). A striking divide in the gene expression pattern was seen for type I vs. type III IFNs by SLRs and poly(I:C) respectively (Fig. 5C). Type III IFNs, IFN-λ2 and IFN-λ3, were more highly expressed upon introduction of poly(I:C) (Fig. 5C). In the mouse, the IFN-λ1 gene is a pseudogene. In addition, matrix metalloproteinases MMP8 and MMP9 (Fig. 5D and SUPP.table 2), which degrade extracellular matrix for cell migration and wound repair, are upregulated to a greater extent by poly(I:C). Similarly, SEMA4F, PLAGL1 and TYRO3 were more upregulated by poly(I:C) injection than SLRs (Fig. 5B and D). Conversely, type I IFNs (10 IFN-α genes and IFN-β) were all elevated by injection of SLRs compared to poly(I:C) (Fig. 5C). In addition to type I IFNs, TNFSF4 (OX40L) and several other genes with known roles in immune function (SATB2, VAV3, and VDR) were more strongly upregulated by SLR14 (Fig. 5 and SUPP.Table 2) (*40*-*42*). A divergent noncoding transcript (9130024F11Rik) that shares a promoter with SATB2 is also highly upregulated (SUPP.Table 2). This suggests a shared mechanism of transcriptional regulation and supports our observations of SATB2 regulation, but the function of these noncoding divergent transcripts is poorly defined.

**Figure 5.**
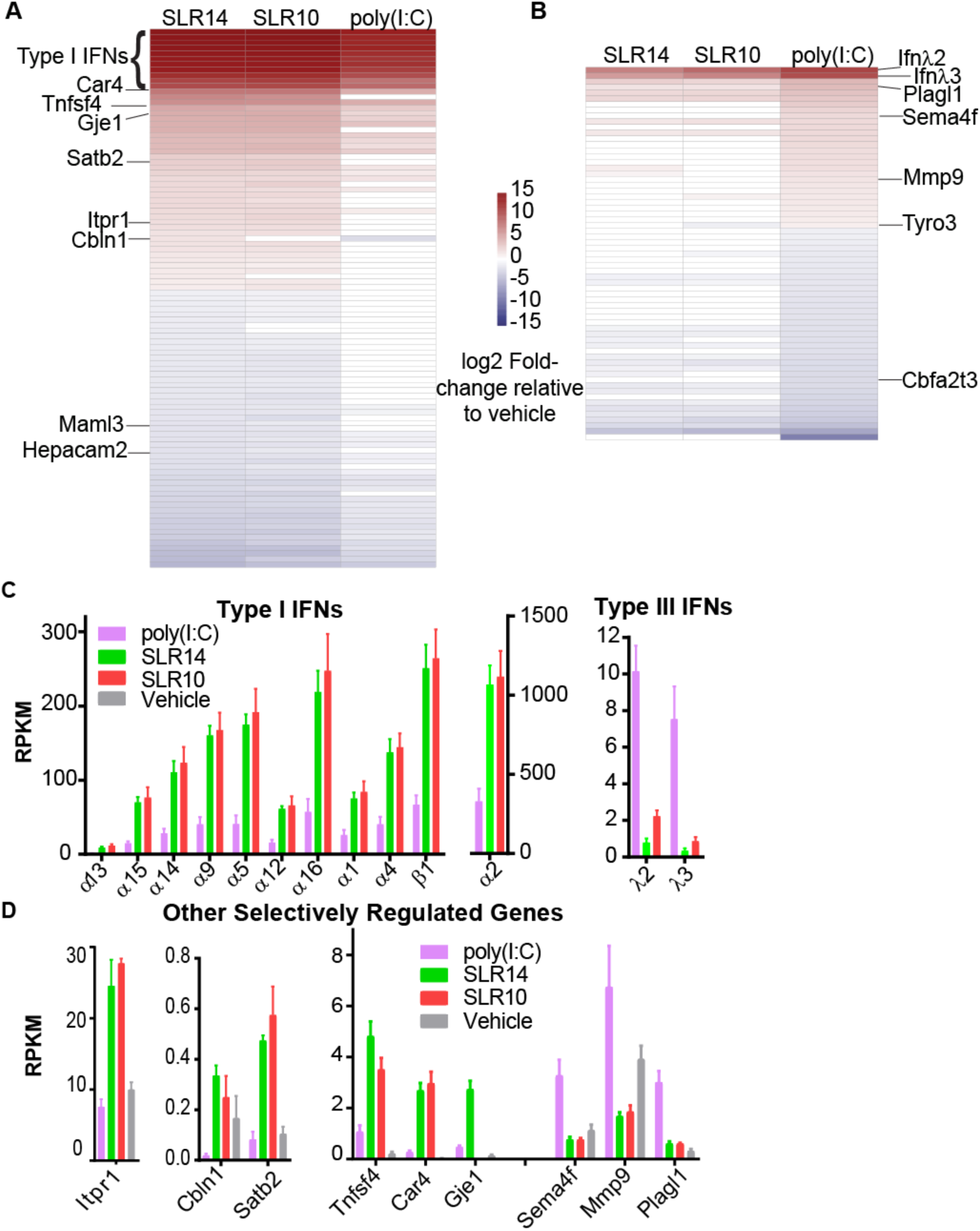
Distinct splenic gene expression between mice treated with SLRs and poly(I:C). Mice were injected i.v. with SLRs or poly(I:C) and 3 hours later, RNA from spleens was purified for RNASeq. Heatmaps were generated for genes preferentially regulated by treatment with SLR (**A**) or poly(I:C) (**B**). The heatmaps include all genes that are differentially expressed between SLR14 and poly(I:C) (> 2 fold-change and FDR < 0.05). Highly statistically significant genes (FDR < 1x10^-5^) are labeled. Type I and type III interferons are preferentially expressed following stimulation with SLR and poly(I:C), respectively (**C**). Several additional genes, with and without reported immune functions, were differentially induced by SLRs or poly(I:C) (FDR < 1x10^-5^) (**D**).

The two treatments also showed differential effects on downregulated genes, including a Rag1/2 repressor (*Zfp608*) that was preferentially downregulated by SLR14 and two repressors of T-cell maturation that were preferentially downregulated by poly (I:C) (*Lax1* and *Dtx1*) (Supp. Table 2) (*43*-*45*). Taken together, these data indicate that SLR14 and related ligands elicit a potent, type I IFN dominant innate immune response while poly(I:C) induces a stronger type III IFN response. Although we have focused on global trends among groups of known genes, the RNAseq data is a rich source of information about potentially novel pathways involved in RNA and antiviral response.

To compare the effects of SLR14 relative to those of triphosphorylated, single-stranded RNA, we examined the DEGs of ppp-NS relative to vehicle, again comparing the response to that of SLR14 (Supp. Figure 4). DEG responses for ppp-NS and SLR14 showed little overlap, confirming that RIG-I is not activated by triphosphate moieties on single-stranded RNA molecules. Taken together, it is now possible to map a network of gene regulatory pathways that are selectively modulated by RIG-I, and which will be activated upon induction of specific RIG-I ligands, such as synthetic, multi-phosphorylated duplexes and viral panhandle RNAs.

## Discussion

RIG-I is a cytosolic sensor that has evolved to detect the presence of RNA molecules that have invaded or been inappropriately expressed within vertebrate cells. Upon binding and responding to these RNA molecules, RIG-I initiates a signaling cascade that leads to pro-inflammatory responses and ultimately apoptosis. In this way, RIG-I plays a key role in the successful innate immune response to viruses, and its activation is being increasingly linked to antitumor response.

Given the antiviral effects of RIG-I, and its potential utility in cancer treatment, it is vital to understand the mechanism of its activation and the molecular structure of its targets, particularly as they are presented in a living animal. This will lead to a better understanding of natural pathways for RIG-I induction, and it will confer the ability to activate and repress RIG-I at will with synthetic RNA molecules and other agonists, giving rise to powerful new therapeutic strategies. Understanding the molecular determinants of RIG-I recognition will also guide the improvement of other therapeutic RNA strategies that elicit unwanted activation of RIG-I, such as siRNA and antisense RNA treatments. We therefore set out to identify a potent, minimal ligand for specific RIG-I induction in animals. Building on crystallographic, biochemical, and cell biological data, we created a set of stable, multiphosphorylated RNA duplexes (the SLRs), and performed comparative testing of their influence on IFN induction and overall gene expression *in vitro* and *in vivo*. By testing these molecules in knockout and knockdown contexts, we have shown that short (≥10 bp), stable RNA duplexes bearing a 5’-triphosphate or diphosphate are potent and specific activators of RIG-I.

The potency of the SLR ligands is significant because biophysical and cell biological experiments from multiple labs has demonstrated that short, polyphosphorylated synthetic RNA duplexes, ranging in length from 10-24 base pairs, can productively bind RIG-I and induce robust RIG-I signaling in cell culture (*3, 9, 10, 22, 24, 26*). These findings challenged the notion that RIG-I forms functional aggregates on RNA, suggesting that RIG-I/MAVS oligomerization is mediated by other features, such as ubiquitination of the CARD domains (*28*-*30*). However, until now, the immune responses and gene expression programs induced by small polyphosphorylated RNA ligands had not been assessed *in vivo*, leaving open the possibility that a functional RIG-I response in an animal requires larger RNA molecules. Here, we establish that polyphosphorylated RNA stemloops as short as ten base-pairs can induce a potent IFN response by RIG-I *in vivo*. This is significant because SLR10 and SLR14 can only bind a single RIG-I molecule, establishing that RIG-I can perform its function without multimerizing on RNA, and validating the *in vivo* activity of short dsRNA. Our findings establish that RIG-I activation can be achieved with stable, synthetic RNA agonists that are chemically well-defined and structurally characterized, which is important for development as a therapeutic. The stemloop design is ideal because it permits RIG-I binding at only a single position on the RNA molecule (the polyphosphate terminus), while the stabilizing UUCG tetraloop enforces a duplex conformation that is resistant to nucleases and strand dissociation. The SLRs therefore represent a powerful new set of tools for exploring the physical basis of RIG-I signaling and for harnessing the therapeutic potential of this process.

The identification of SLR agonists provides a unique opportunity to explore the gene expression pathways that are specifically controlled by RIG-I, and to differentiate them from networks that are initiated by other types of receptors. This has not been possible in the past because RNAs used in previous studies of RIG-I response (such as poly(I:C) or Sendai virus RNA) have large, complex structures that are capable of recruiting many types of receptors (such as MDA5, TLR3 and TLR7), and binding a host of potential coactivator proteins. Furthermore, the majority of studies of RIG-I induction have been conducted in cell culture systems, and there is a lack of transcriptome-wide analysis of RIG-I induction in living animals. We therefore utilized the SLRs as tools to examine gene expression upon RIG-I activation *in vivo*, and to compare the SLRs with other types of RNA molecules.

As expected, SLRs and poly(I:C) both induce a large number of genes that are associated with antiviral immunity and ISGs. However, SLRs selectively induced elevated levels of genes belonging to type I IFNs, whereas poly(I:C) induced higher levels of type III IFN genes (IFN-λ2 and IFN-λ3). While the precise mechanisms by which a given RIG-I agonist triggers type I vs. type III IFN genes is unknown, recent studies highlight that the intracellular location of MAVS might dictate these differential gene activation pathways. Kagan and colleagues have shown that MAVS residing in the peroxisomes preferentially induce type III IFN expression, while MAVS residing on the surface of mitochondria activate the transcription of type I IFN (*46*). Both of these pathways require IRF3 to induce the IFN genes, but the type III pathway depends on IRF1, while the type I pathway depends on Erk1. Therefore, it is conceivable that MAVS activation is preferentially induced at the mitochondria by SLRs, and at the peroxisome by poly(I:C). An alternative hypothesis is that SLRs trigger a specific cell type that preferentially induces type I IFNs, while poly(I:C) stimulates another cell type that selectively induces type III IFNs. Future studies are needed to distinguish these possibilities by analyzing the cell types responsible for the secretion of respective IFNs *in vivo*. Whatever the mechanism might be, our comprehensive transcriptome results have important implications for the use of RNA ligands in humans. For instance, type I IFNs play a key role in initiating anti-tumor CD8 T cell responses (*47*-*49*). Thus, SLRs are expected to better elicit type I IFNs during cancer immunotherapy. On the other hand, systemic recombinant IFN-α therapy using conventional methods is not well tolerated due to significant side effects, so it may be prudent to minimize the type I response, particularly for localized pathologies. For example, hepatocytes express IFN-λR during the antiviral response against HCV and might benefit from a type III IFN response (*50, 51*). Thus, poly(I:C) may be a more effective therapy for HCV given its preferential induction of type III IFNs.

Our RNAseq analysis revealed RIG-I induction of unexpected gene families, including several without annotated immune functions, suggesting potentially novel pathways involved in RIG-I response. For example, genes involved in pH homeostasis (*Car4* and *Sct*) were selectively upregulated by SLR14. Local acidosis has long been associated with inflammation, and these genes may be indicative of a pathway that functions as a response to this environment (*52*-*54*). Furthermore, the maintenance of pH homeostasis could benefit an immune response to tumors with acidic microenvironemnts. Three other genes preferentially upregulated by SLR14 have been primarily characterized as neuronal proteins (*Gje1, Cbln1*, and *Itpr1*). Further investigation of these neuronal proteins revealed homology with known effectors of immune function (connexins, complement domains, and inositol receptors, respectively), which may suggest a *bone fide* role for these genes in the immune response (*55*-*58*).

RIG-I induction and antiviral responses are expected to cause restructuring of cellular structure and metabolism, and it is therefore intriguing that specific sets of gene families are downregulated upon systemic RIG-I activation in the animal. For example, membrane proteins are significantly downregulated upon stimulation. In particular, genes involved in cell-cell contacts are downregulated, which is consistent with the migration of cells out of the spleen to peripheral tissues upon immune activation. Furthermore, C-type lectins are downregulated, including many expressed on dendritic, macrophage, and NK cells, suggesting a depletion of these cells in the spleen as they migrate to peripheral tissues (*59, 60*).

In summary, we have demonstrated that SLRs are highly potent activators of the IFN response in living animals. They provide valuable tools for exploring the molecular determinants of RIG-I activation in the complex environment of the mammalian immune system and they facilitate characterization of the gene expression pathways that are controlled by the RIG-I receptor. Given their small size and chemically-defined composition, SLRs also represent a promising new class of oligonucleotide therapeutics with potential applicability as antivirals, vaccine adjuvants and antitumor agents.

### Materials and Methods

#### RNA synthesis and purification

All triphosphorylated RNA oligonucleotides were synthesized on a MerMade 12 DNA-RNA synthesizer (BioAutomation), using previously described synthetic procedure (*61, 62*). The 2’-pivaloyloxymethyl phosphoramidites were obtained from ChemGenes. Synthesized triphosphorylated RNAs were deprotected with ammonium hydroxide as described (*62*) and purified by polyacrylamide gel electrophoresis. RNA molecules were further analyzed for purity by mass spectrometry (Novatia LLC). Low molecular weight poly(I:C) was purchased from Invivogen and used without further purification.

#### HEK 293T Cell Culture and IFN-β Induction Assays

Cell based experiments were conducted in HEK 293T cells because they do not express endogenous RLRs (proteinatlas.org). Cells were grown and maintained in 15 cm dishes containing Dulbecco’s Modified Eagle Medium (DMEM; Life Technologies) supplemented with 10% heat-inactivated fetal bovine serum and Non-Essential Amino Acids (Life Technologies). IFN-β induction assays were conducted in 24-well format. Briefly, 0.5 mL of cells at 100,000 cells/mL were seeded in each well of tissue culture treated 24-well plates. After 24 hours, each well of cells was transfected with 3 ng of pUNO-hRIG-I or pUNO-hMDA5 (Invivogen), 6 ng pRL-TK constitutive Renilla luciferase reporter plasmid (Promega), and 150 ng of an IFN-β/Firefly Luciferase reporter plasmid using the Lipofectamine 2000 transfection reagent (Life Technologies) per the manufacturer’s protocol. Protein expression was allowed to proceed for 24 hours, at which point the cells were challenged by transfection of 1 μg of the indicated RNA or LMW polyI:C (6 wells per RNA condition), also using the Lipofectamine 2000 reagent. After 12 hours, cells were harvested for luminescence analysis. These experiments were performed in biological triplicate.

Induction of the IFN-β promoter was analyzed using a dual luciferase assay. After aspiration of the growth media, 100 μL of passive lysis buffer (Promega) was added to each well, and incubated for 15 min at room temperature. The lysed cells were transferred to a 96-well PCR plate and clarified by centrifugation. Next, 20 μL samples of the supernatant were transferred to a 96-well assay plate for analysis using the Dual-Luciferase Reporter Assay System (Promega). Luminescence was measured using a Biotek Synergy H1 plate reader. The resulting Firefly luciferase activity (i.e. the induction of IFN-β) was normalized to the activity of the constitutively expressed Renilla luciferase to account for differences in confluency, viability and transfection efficiency across sample wells.

#### Mice

C57BL/6NCrl (WT) mice were purchased from Charles River Laboratories (Frederick, MD). *Tlr7* (The Jackson Laboratory; B6.129S1-Tlr7tm1Flv/J) and *Mavs* knockout mice (The Jackson Laboratory; B6;129-Mavstm1Zjc/J) were backcrossed to C57BL/6 mice for several generations. *Mda5* knockout mice (The Jackson Laboratory; B6.Cg-Ifih1tm1.1Cln/J) were a kind gift from Dr. Erol Fikrig. All animal procedures were performed in compliance with Yale Institutional Animal Care and Use Committee protocols.

#### In vivo injection of SLRs

SLRs were complexed to *in vivo*-jetPEI^®^ (Polyplus transfection) according to manufacturer’s instructions, with N/P ratio = 8. Unless otherwise indicated, 25 μg of RNA in a 200 μl volume was injection per mouse intravenously. At various time points, sera were collect and ELISA (PBL Assay Science, eBioscience) was performed according to manufacturers instructions. Spleens were collected into RNAlater (QIAGEN) for RNA extraction. A minimum of 3 mice per group was used per experiment.

#### RNA extraction from spleen

Tissues collected into RNAlater were blotted dry and transferred into Trizol (Thermo Fisher Scientific) in a Lysing Matrix D tube (MP Biomedical) for homogenization. Supernatant was collected for chloroform phase separation and RNA was precipitated using isopropyl alcohol. The RNA pellet was then washed with 75% ethanol and redissolved in water.

#### Quantification of gene expression in animals by qRT-PCR

cDNA was synthesized from RNA using iScript cDNA synthesis kit (Bio-Rad). Quantitative PCR was performed using iTaq Universal SYBR Green (Bio-Rad) on a CFX Connect instrument (Bio-Rad). *Ifna-4* (F: CTG CTA CTT GGA ATG CAA CTC; R: CAG TCT TGC CAG CAA GTT GG) and *Mx1* (F: TGT ACC CCA GCA AAA CAT CA; R: TTG GAA GCG CTA AAG TGG AA) expression was normalized to *Hprt* (F; GTT GGA TAC AGG CCA GAC TTT GTT G; R: GAG GGT AGG CTG GCC TAT TGG CT).

#### RIG-I knockdown experiments

A549 cells (ATCC, Manassas, Va., CCL-185) were propagated in F-12K Medium (Kaighn’s Modification of Ham’s F-12 Medium) containing 10% fetal bovine serum (FBS). Cells were reverse transfected in a 24-well plate with a RIG-I (DDX58)-specific siRNA sequence (GE Healthcare Dharmacon Inc Cat# D-012511-01-005) at a final concentration of 25 nM RNA and 0.5 μl lipofectamine 2000 (Thermo Fisher Scientific) per well, according to the manufacturer’s instructions. After 4h incubation at 37 °C, transfection medium was replaced with fresh growth medium and incubated for another 20h at 37 °C. Following siRNA treatment, cells were challenged with either mock (transfection reagent only) or 1.7 ng/μl exogenous purified RNA (SLR14 or SLR10) and μl of Lipofectamine2000 reagent (Thermo Fisher Scientific) per well, according to the manufacturer’s instructions. Cells were transfected for 6h at 37**°**C, then trypsinized, washed in phosphate-buffered saline (PBS), and pelleted for total RNA extraction.

Total RNA from cells was extracted using the RNAeasy Mini Kit (Qiagen, Valencia, CA, USA) protocol. Genomic DNA contamination was removed by DNAseI treatment (Qiagen) directly on the Mini kit column. Total RNA was reverse transcribed into cDNA using oligo dT primers and Superscript III reverse-transcriptase enzyme (Thermo Fisher Scientific) according to the manufacturer’s instructions. Quantitative real-time PCR was performed using CFX384 Touch™ Real-Time PCR Detection System (BioRad) and LightCycler^®^ 480 SYBR Green I Master (Roche Diagnostics). Each sample was analyzed in technical and biological triplicate and normalized to HPRT expression. Gene expression quantification was performed according to the ΔΔCt method and fold change values are reported relative to cells without siRNA treatment and mock challenged with RNA. Primer sequences are: Hprt (F: TTTCAAATCCAACAAAGTCTGGC; R: TGGTCAGGCAGTATAATCCAAAG), Rig-I (F: TTCATGTCCACCTTCAGAAGTG; R: TCATAGCAGGCAAAGCAAGC), Ifnb1 (F:GTCACTGTGCCTGGACCATAG; R: GTTTCGGAGGTAACCTGTAAGTC), Rsad2 (F: TCGCTATCTCCTGTGACAGC; R: CACCACCTCCTCAGCTTTTG), Mx1 (F: AGAGAAGGTGAGAAGCTGATCC; R: TTCTTCCAGCTCCTTCTCTCTG)

#### RNA seq library preparation and data analysis

Libraries were prepared using Illumina TruSeq Stranded mRNA sample preparation kits from 500ng of purified total RNA according to the manufacturer’s protocol. The finished dsDNA libraries were quantified by Qubit fluorometer, Agilent TapeStation 2200, and RT-qPCR using the Kapa Biosystems library quantification kit according to manufacturer’s protocols. Uniquely indexed libraries were pooled in equimolar ratios and sequenced on an Illumina NextSeq500 with paired-end 75bp reads by the Dana-Farber Cancer Institute Molecular Biology Core Facilities.

#### Bioinformatic Analysis

Sequenced reads were checked for quality control using the FastQC tool (Babraham Bioinformatics, Babraham Institute, Cambridge, UK. Good quality reads were mapped to the reference genome (GRCm38/mm10) using the software package Tophat v 2.0.6 (*63*). Default settings were used, except for the mean inner mate distance between the pairs that was set to 150bp. A maximum of two mismatches and a minimum length of 36 bp per segment were allowed. The BAM files from Tophat were then converted to SAM format by Samtools (v 1.4) (*64*) and raw counts estimated by the Python script HTSeq count (*65*). Reference GTF file for gene annotation was downloaded from iGenomes website (http://cufflinks.cbcb.umd.edu/igenomes.html).

We generated approximately 628 million paired-end sequencing reads by multiplexing 24 samples on 4 lanes of Illumina NextSeq500 platform. On average, each sample achieved a sequencing depth of 26 million paired-end reads. The overall mapping rate against the reference transcriptome up to 92%. Normalized gene expression levels, represented as counts per million (CPM), were calculated for 39,169 transcripts from the Ensembl database and were processed for statistical analysis with EdgeR (*66*).

The resulting raw counts per gene were processed by the EdgeR program to perform normalization, clustering and estimate differential expression. EdgeR (Bioconductor release 3.4.2) performs differential abundance analysis using a pair-wise design based on negative binomial model, as an alternative to the Poisson estimates. Normalization of the sequenced libraries was performed to remove effects due to differences in library size. The resulting p-values were corrected for multiple testing using the Benjamini-Hochberg false discovery rate (FDR) approach.

We identified genes uniquely regulated by SLR14 or poly(I:C) using a two-step filtration process. First, each treatment was compared to vehicle and differentially expressed genes were selected that had p-values <0.05 and >2-fold change in expression level. This subset of genes was then compared between the SLR14 and poly(I:C) treatments. Genes that were differentially expressed between SLR14 and poly(I:C) with p-values <0.05 and >2-fold change in expression level were deemed “significantly more up/downregulated by a specific treatment”. All other genes were deemed “mutually up/downregulated”. Mutually up/downregulated genes were analyzed for functional enrichment using the Database for Annotation, Visualization and Integrated Discovery (DAVID) v6.8 (*67*).

## Acknowledgments

### Funding

This study was supported in part by the Howard Hughes Medical Institute, and awards from the National Institutes of Health (R01AI054359 and R56AI125504 to A.I.)

#### Competing interests

SLRs are the subject of patent number 14/776,463

## Supplementary Materials

### RNA Characterization

SLRs are palindromic sequences terminated by a stabilizing UUCG tetraloop, and are therefore most stable as monomeric stemloop (Fig. 1A). Although the monomeric nature of the SLRs has been confirmed by crystallography, ultracentrifugation (*10*), and fluorescence anisotropy experiments (*38*), at high SLR concentrations, we used native-gel electrophoresis to examine their propensity to form self-complementary duplexes (dimers) and to assess the purity of the monomeric stemloop form under the conditions used in this study (including the lipofectamine formulation). Non-palindromic triphosphorylated oligonucleotides of identical internal duplex stability were synthesized as dimerization controls. Under all conditions SLRs migrate exclusively as monomeric hairpin RNA molecules, without any trace of dimerized product (Supplementary Fig. 1), confirming that SLR RNA is monomeric when it is introduced into cells and animals. We cannot rule out the possibility of disproportionation once SLRs enter cells, but given the increased dilution upon transfection and the high entropic cost of dimerization, it is highly unlikely.

**Supplementary Figure 1:**
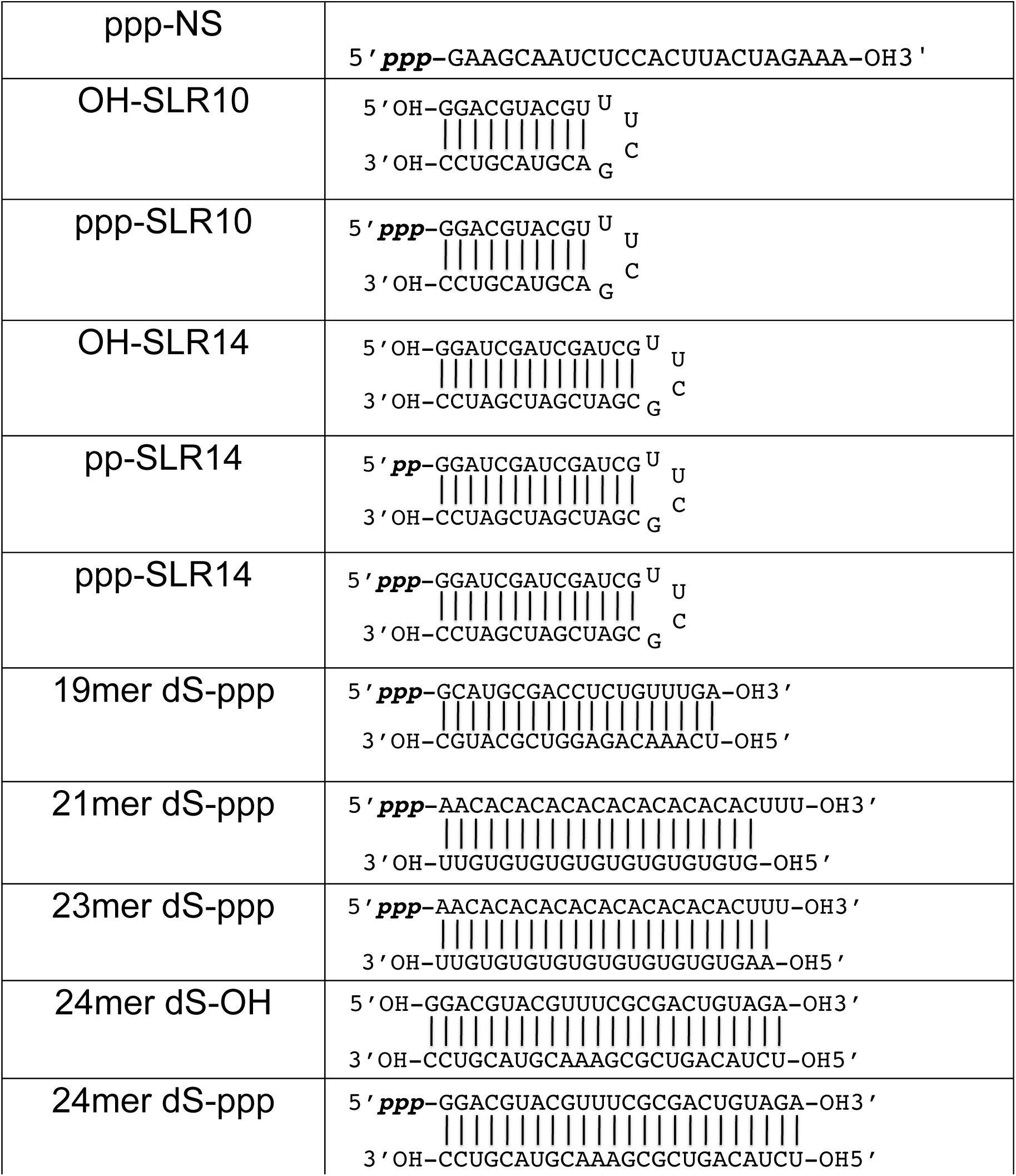
RNA ligands for RIG-I activation. All oligonucleotides were synthesized as described in Methods. The dsRNAs used in this experiment vary in length and stability, and were tested to facilitate comparisons with published studies on annealed duplexes as RIG-I ligands. The 24mer dsRNA duplex (43.9 kcal/mol stability) has a terminal sequence analogous to SLR10 and SLR14, and approximately twice the duplex length. The 19mer dsRNA duplex (32.7 kcal/mol) is the same sequence and reported composition as an RNA that is marketed as a RIG-I ligand (Invivogen), but it was synthesized in-house, in parallel with all other RNAs used in this study. The 21mer dsRNA (35.9 kcal/mol) & 23mer dsRNA (37.4 kcal/mol) duplexes are identical in sequence and reported composition to those reported previously in studies of RIG-I activation (*9*). Importantly, these have a variable 3’-overhang on the non-triphosphorylated duplex end. Free energies of all duplexes, including overhanging ends, were calculated as described (*68*).

**Supplementary Figure 2.**
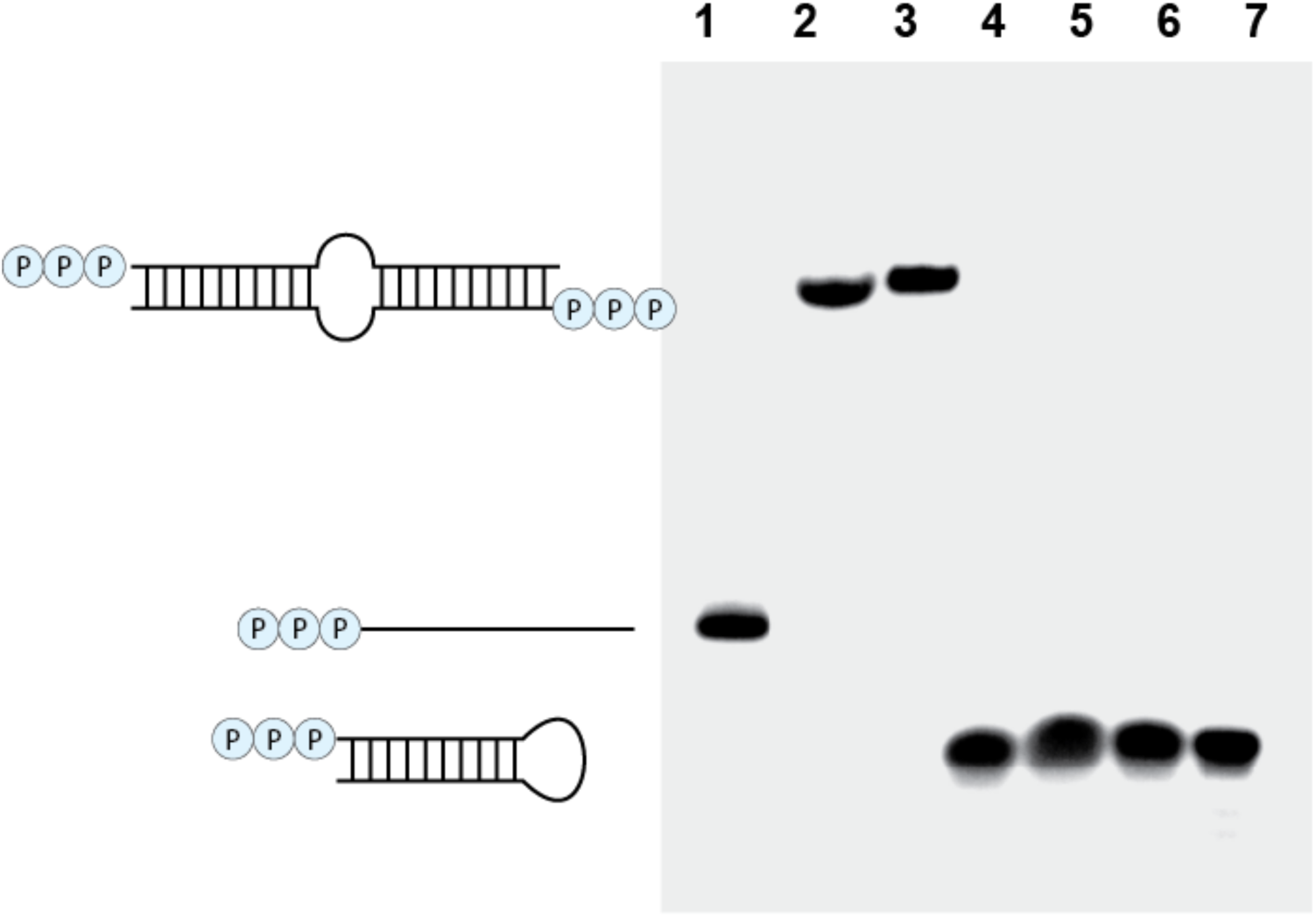
SLR10 is monomeric in conditions that mimic cell culture and *in vivo* experiments. Monomeric and dimeric RNA species are separated on a native gel. **1)** 24-mer single stranded RNA marker (5’ - pppUCUACAGUCGUUCGACGUACGUCC - 3’); **2)** 24-mer duplex marker corresponding to a putative SLR10 dimer control (5’ - pppGGACGUACGUUUCGCGACUGUAGA – 3’ / 5’ – pppUCUACAGUCGUUCGACGUGCAUCC – 3’). 0.5 mM RNA is in ME buffer (10 mM Na-MOPS pH 6.0, 1 mM trisodium EDTA); **3)** 24-mer duplex marker in cell culture reagents: 0.001 mM RNA in Opti-MEM + lipofectamine; **4)** 0.5 mM SLR10 in ME buffer, heated to 95 °C and snap cooled immediately prior to running to control for freezing/thawing effect on HP folding; **5)** 1 mM SLR10 in ME buffer, thawed from −80 °C and loaded without refolding; **6)** 0.5 mM SLR10 in ME buffer, thawed from −80 °C; **7)** SLR10 in cell culture reagents: 0.001 mM RNA in Opti-MEM + lipofectamine

**Supplementary Figure 3.**
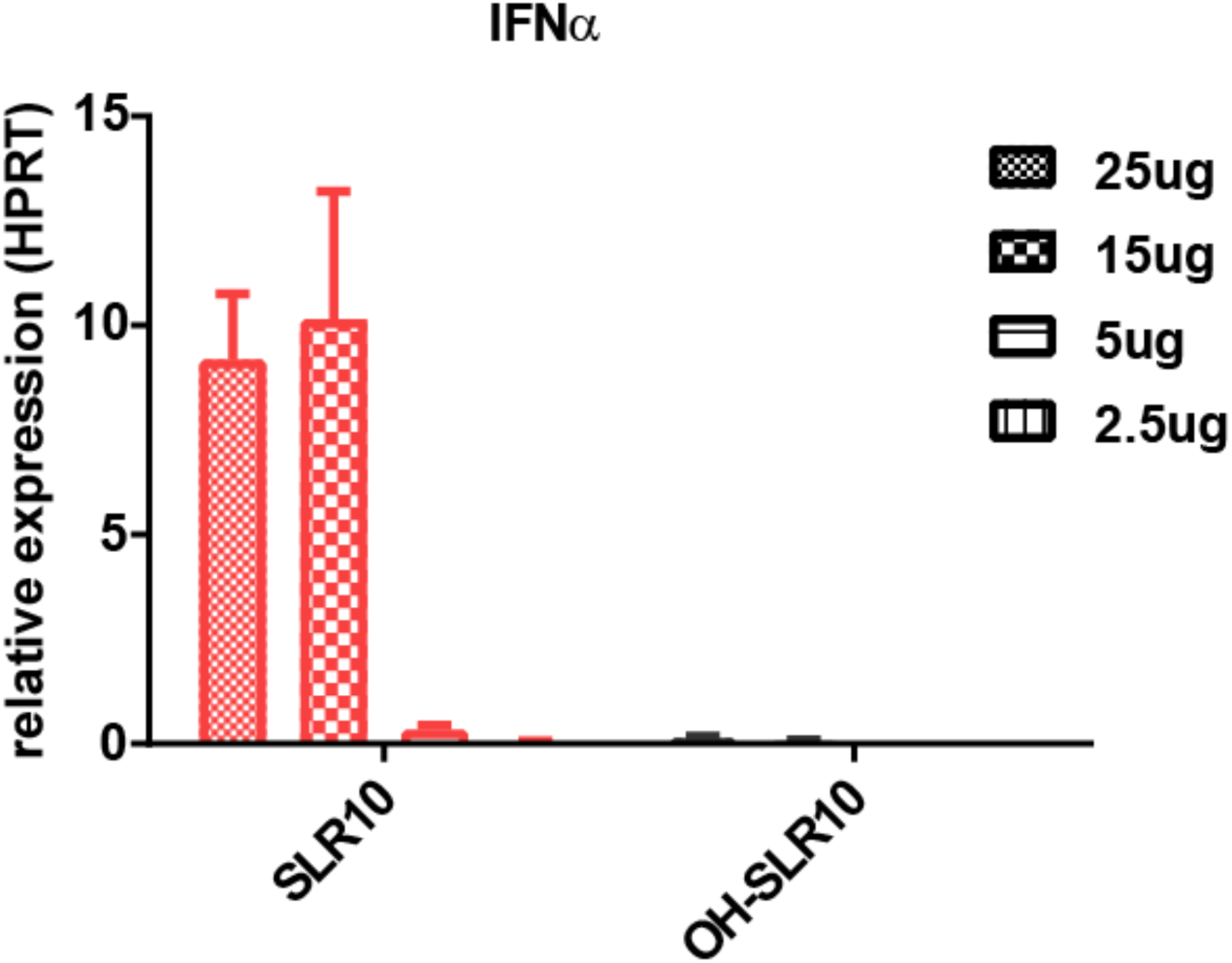
Dose response of SLRs. Mice were injected i.v. with SLR10 or OH-SLR10 and spleens harvested after 3 hours for RNA isolation. IFNα was measured by RT-qPCR. 25ug was chosen as the optimal dose for all subsequent experiment.

**Supplementary Figure 4.**
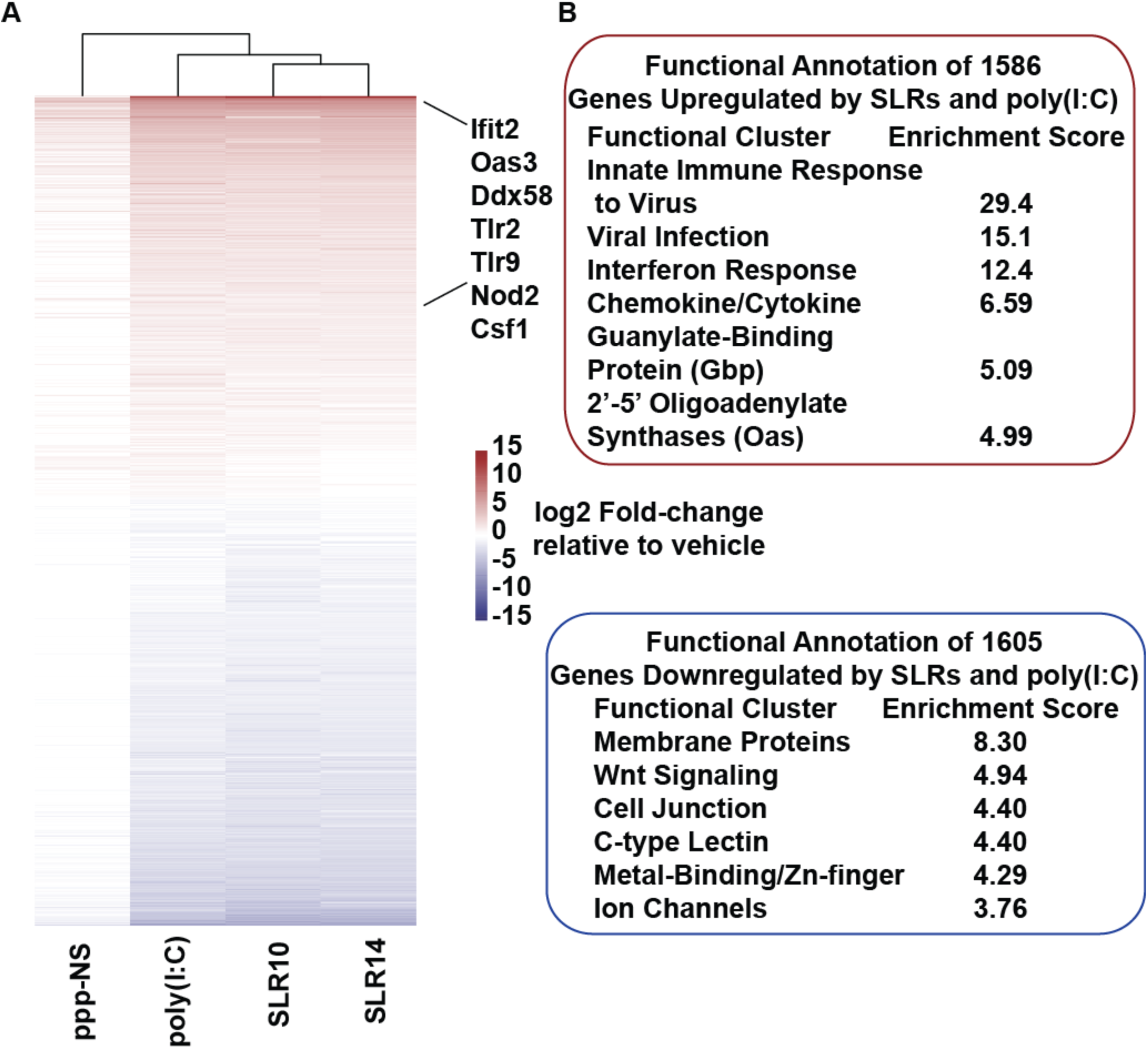
SLRs and poly(I:C) regulate a shared set of genes. Heatmaps were generated for all genes that were differentially expressed relative to vehicle upon treatment with RNA (**A**). Many genes associated with innate immunity and antiviral response were upregulated by SLR and poly(I:C) to similar extents. Several examples are indicated to the right of the heatmap. DAVID pathway analysis was performed on the annotated functions of genes mutually up/downregulated by treatment with SLR and poly(I:C) (**B**). Genes selected for analysis were differentially expressed by SLR14 or poly(I:C) relative to vehicle (>2 fold-change, FDR < 0.05) but were not differentially expressed between the two treatments (< 2 fold-change or FDR > 0.05).

**Data file S1. List of genes mutually regulated by SLRs and poly(I:C).** A list and statistical metrics for all genes that are differentially expressed by SLR14 or poly(I:C) relative to vehicle (>2 fold-change and FDR < 0.05) but not differentially expressed between SLR and poly(I:C) (< 2 fold-change or FDR > 0.05). Complete results from the DAVID analysis in Supplementary Figure 4B are also included.

**Data file S2. List of genes preferentially regulated by SLRs or poly(I:C).** A list and statistical metrics for all genes that are differentially expressed by SLR14 or poly(I:C) relative to vehicle (>2 fold-change and FDR < 0.05) and differentially expressed between SLR and poly(I:C) (> 2 fold-change and FDR < 0.05). These data correspond to Figure 5.

